# The multi-scale architecture of mammalian sperm flagella and implications for ciliary motility

**DOI:** 10.1101/2020.11.18.388975

**Authors:** Miguel Ricardo Leung, Marc C. Roelofs, Ravi Teja Ravi, Paula Maitan, Min Zhang, Heiko Henning, Elizabeth G. Bromfield, Stuart C. Howes, Bart M. Gadella, Hermes Bloomfield-Gadêlha, Tzviya Zeev-Ben-Mordehai

## Abstract

Motile cilia are molecular machines used by a myriad of eukaryotic cells to swim through fluid environments. However, available molecular structures represent only a handful of cell types, limiting our understanding of how cilia are modified to support motility in diverse media. Here, we use cryo-focused ion beam milling-enabled cryo-electron tomography to image sperm flagella from three mammalian species. We resolve in-cell structures of centrioles, axonemal doublets, central pair apparatus, and endpiece singlets, revealing novel protofilament-bridging microtubule inner proteins throughout the flagellum. We present native structures of the flagellar base, which is crucial for shaping the flagellar beat. We show that outer dense fibers are directly coupled to microtubule doublets in the principal piece but not in the midpiece. Thus, mammalian sperm flagella are ornamented across scales, from protofilament-bracing structures reinforcing microtubules at the nano-scale to accessory structures that impose micron-scale asymmetries on the entire assembly. Our structures provide vital foundations for linking molecular structure to ciliary motility and evolution.

## Introduction

Cilia, also called flagella, are evolutionarily ancient organelles used by a menagerie of eukaryotic cell types and organisms to propel themselves through fluid environments (Mitchell, 2017; Wan, 2018). These intricate molecular machines are paragons of self-organization built from a bewildering array of active and passive structural elements that, together, are able to spontaneously generate oscillatory wave-like motion (Gaffney et al., 2011). The basic architecture of motile cilia is conserved across broad swaths of the eukaryotic tree, providing information on the minimal structures needed for spontaneous undulation (Brokaw, 2009). However, because they operate in a wide range of environments, cilia from different cell types generate different waveforms (Khan and Scholey, 2018) that are further modulated by fluid viscosity (Smith et al., 2009).

The motile cilium is a continuous assembly of compound microtubules (Ishikawa, 2017). The base of the cilium is the centriole or basal body, which is typically a cylinder of triplet microtubules. The centriole transitions into the axoneme, which consists of nine doublet microtubules arrayed around a central pair of singlet microtubules. Axonemal microtubules anchor hundreds of dynein motors and accessory proteins to power and regulate movement. Axoneme structure has been studied extensively by cryo-electron tomography (cryo-ET) in *Chlamydomonas*, *Tetrahymena*, and sea urchin sperm (Nicastro et al., 2006, 2011; Owa et al., 2019; Pigino et al., 2012). Recent studies have begun to shed light on species- and cell type-specific specializations (Greenan et al., 2020; Imhof et al., 2019; Lin et al., 2014a; Yamaguchi et al., 2018), motivating efforts to expand the pantheon of organisms and cell types used in axoneme research.

Perhaps the most striking example of ciliary diversity across species is in sperm, which are highly specialized for a defined function – to find and fuse with the egg. Sperm consist of a head, which contains the genetic payload, and a tail, which is a modified motile cilium. Despite their streamlined structure, sperm are simultaneously the most diverse eukaryotic cell type (Gage, 2012; Lüpold and Pitnick, 2018), reflecting the sheer range of reproductive modes and fertilization arenas, from watery media for marine invertebrates and freshwater species to the viscous fluids of the female reproductive tract for mammals. Because motility is crucial to sperm function, the natural variation of sperm form thus presents a unique opportunity to understand the structural diversification of motile cilia.

Mammalian sperm flagella are characterized by accessory structures that surround and dwarf the axoneme (Fawcett, 1975), unlike marine invertebrates whose sperm tails consist essentially of the axoneme (Fawcett, 1970). In mammalian sperm, axonemal doublets are associated with filamentous cytoskeletal elements called outer dense fibers (ODFs) for most of their lengths. The ODFs are further surrounded by a sheath of mitochondria in the midpiece and by a reticular structure called the fibrous sheath in the principal piece. These accessory structures are proposed to stabilize beating of the long flagella of mammalian sperm (Lindemann, 1996; Lindemann and Lesich, 2016). The accessory structures may also facilitate movement through viscous media by suppressing buckling instabilities that would otherwise cause sperm to swim in circles (Gadêlha and Gaffney, 2019). Indeed, many cases of male infertility are linked to defects in these accessory elements (Haidl et al., 1991; Serres et al., 1986; Zhao et al., 2018). However, we still do not fully understand how these accessory structures modulate the flagellar beat since there is very little structural information on how they interact with the axoneme proper.

Another distinguishing feature of mammalian sperm flagella is that they are not anchored by a basal body (Avidor-Reiss, 2018). Instead, the base of the mammalian sperm flagellum is surrounded by a large cytoskeletal scaffold called the connecting piece. The isolated bovine sperm connecting piece was characterized by cryo-ET, revealing a complex asymmetric assembly (Ounjai et al., 2012). However, the purification process resulted in loss of the centrioles. Thus, there is still a paucity of structural information on the flagellar base in intact cells and on how it varies across species that often have very different head shapes.

Sperm have two centrioles that are located in the neck, where the nucleus attaches to the flagellum. The centriole closer to the nucleus is referred to as the proximal centriole (PC) and the one at the base of the flagellum the distal centriole (DC). During spermiogenesis in mammals, the DC is remodeled to the point that it no longer resembles a canonical centriole. This was thought to represent a process of degeneration (Manandhar et al., 2000), but recent work showed that the DC is in fact a functional centriole that participates in orchestrating the first zygotic division (Fishman et al., 2018). Such drastic deviations from canonical centriole architecture have not been investigated in detail.

Here, we combine cryo-focused ion beam (cryo-FIB) milling-enabled cryo-ET (Marko et al., 2007; Rigort et al., 2012) with subtomogram averaging to image mature sperm from three mammalian species – the pig (*Sus scrofa*), the horse (*Equus caballus*), and the mouse (*Mus musculus*) – that differ in terms of gross morphology, motility, and metabolism. We leverage the uniquely multiscale capabilities of cryo-ET to define the molecular architecture of microtubule-based assemblies and their critical interactions with accessory structures. We take advantage of the highly streamlined shape of sperm in order to define how these structures and relationships change throughout the flagellum.

We define the architecture of the flagellar base and show that ODFs are anchored through a large, asymmetric chamber around the centrioles. We show that ODFs are directly coupled to axonemal microtubules in the principal piece, but not in the midpiece. We find that mammalian sperm microtubules are additionally decorated throughout by protofilament-bridging microtubule inner protein densities. Thus, mammalian sperm flagella are modified across scales – from large accessory structures that increase the effective size and rigidity of the entire assembly to extensive microtubule inner proteins that likely reinforce the microtubules themselves. We further discuss the implications of this multi-scale architecture for ciliary motility.

## Results

### The base of the flagellum is anchored through a large, asymmetric chamber around the centrioles

The neck region containing the PC and DC is too thick (~600-700 nm) for direct imaging by cryo-ET, so in order to image sperm centrioles in their native subcellular milieu, we used cryo-FIB milling to generate thin lamellae suitable for high-resolution imaging **(Fig. 1)**. Cryo-ET of lamellae containing the PC confirmed that it is indeed composed of triplet microtubules in pig and in horse sperm **(Fig. 1a-b)**. Unexpectedly, we found that triplets of the pig sperm PC are not all the same length **(Fig. 1a, S1a)**. Shorter triplets are grouped on one side of the centriole, giving the PC a striking dorsoventral asymmetry **(Fig. S1a)**. Consistent with previous reports that the PC degenerates in rodents (Manandhar et al., 1998; Woolley and Fawcett, 1973), the PC was not prominent in mouse sperm. However, cryo-ET showed unequivocally that some centriolar microtubules remain **(Fig. 1c)**, demonstrating that degeneration is incomplete. We observed complete triplets as well as triplets in various stages of degeneration, including triplets in which only the B-tubule had degraded **(Fig. 1c’)**.

**Fig. 1.**
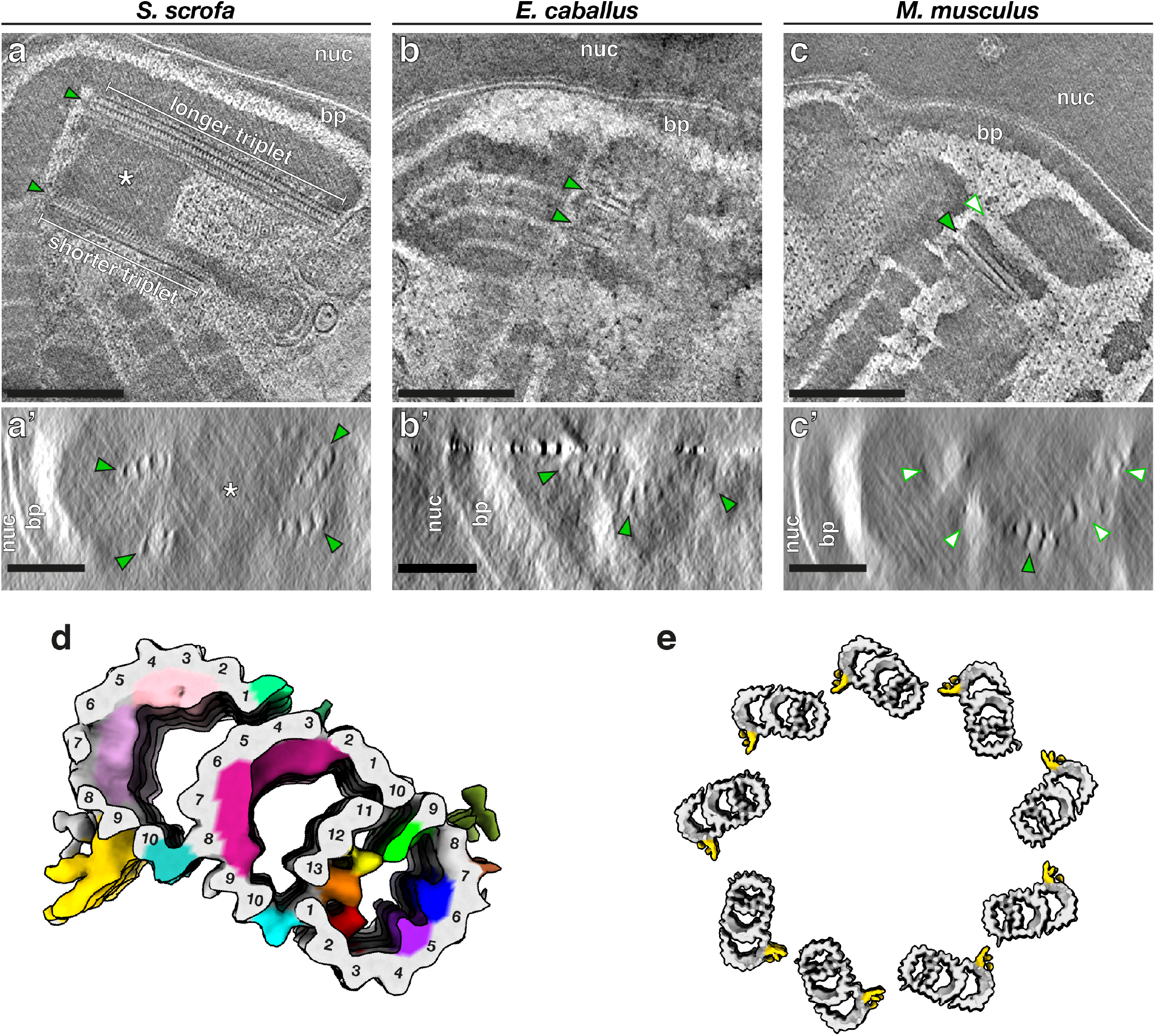
The proximal centriole (PC) in mammalian sperm is asymmetric and contains novel microtubule inner proteins. Tomographic slices through cryo-FIB milled lamellae of pig **(a)**, horse **(b)**, and mouse **(c)** sperm. Transverse slices **(a’-c’)** show complete triplets in the pig **(a’)** and the horse **(b’)**, but not in the mouse **(c’)**. Complete triplets are indicated by green arrowheads with black outlines, while degenerated triplets are indicated by white arrowheads with green outlines. **(d)***In situ* structure of the pig sperm PC with the tubulin backbone in grey and microtubule inner protein densities colored individually. **(e)** Reconstruction of the pig sperm PC with the A-C linker colored in yellow. **Labels:** nuc - nucleus, bp - baseplate. **Scale bars:**(a,b,c) 250 nm; (a’,b’,c’) 100 nm.

We determined the *in situ* structure of the pig sperm PC by subtomogram averaging **(Fig. 1d)**. While the overall structure of the PC triplet is similar to other centriole structures (Greenan et al., 2018, 2020; Guennec et al., 2020; Guichard et al., 2013; Li et al., 2012), it differs in terms of the microtubule inner protein densities (MIPs) **(Fig. S1b)**. We observed nine MIPs, six in the A-tubule, one in the B-tubule, and two in the C-tubule (Fig. **(Fig. S1c)**. In the A-tubule most of the MIPs are unique, including MIP2 (yellow) that binds to protofilament A12, MIP3 (orange) bridging protofilaments A13 and A1, MIP4 (red) that binds to A2, MIP5 (purple) that binds to A5, and MIP6 (blue) bridging A6 and A7. MIP1 (green), a prominent MIP associated with protofilament A9, was also reported in centrioles isolated from CHO cells (Greenan et al., 2018) and in basal bodies from bovine respiratory epithelia (Greenan et al., 2020). Protofilaments A9 and A10 are proposed to be the location of the seam (Ichikawa et al., 2017), which suggests that MIP1 is a highly-conserved seam-stabilizing or seam-recognizing structure.

In the B-tubule, we observed a large helical MIP7 (magenta) bridging protofilaments B3-9. We observed two groups of unique MIPs in the C-tubule, MIP8 (light pink) associated with C2-C4 and MIP9 (pink) with C5-C7. The inner junctions between A- and B-tubules (cyan) and between B- and C-tubules (turquoise) are non-tubulin proteins that repeat every 8 nm and are staggered relative to each other when viewed from the luminal side of the triplet. We resolved density for the A-C linker (gold), which is associated with protofilaments C9 and C10 **(Fig. 1e)**.

The B- and C-tubule MIPs we observed are not present in other centriole structures. However, helical MIPs have been observed in the transition zone of bovine respiratory cilia (Greenan et al., 2020). Unlike other mammalian centriole structures, we do not observe MIPs that bridge B1-B2 or C1-C2 **(Fig. S1b)**. It is difficult to tell whether these differences in MIP patterns are due to differences in cell type or species. As this is the first *in situ* structure of any mammalian centriole, these differences may also be because previous structures were of isolated centrioles. Nonetheless, it is clear that there is great diversity in how core centriolar microtubules are accessorized, which raises questions about the functions of these MIPs.

We next determined the organization of the atypical DC by tracing microtubules through Volta phase plate (VPP) (Danev et al., 2014) cryo-tomograms of whole sperm **(Fig. 2)**. The DC consists of doublet microtubules, with a pair of singlets extending through the lumen **(Fig. 2a-f)**. In pig and in horse sperm **(Fig. 2a-d)**, doublets extend almost as far proximally as the central pair. In a further departure from canonical centriole structure, DC doublets are splayed open and arranged asymmetrically around the singlets. The central singlets themselves are spaced inconsistently, suggesting that they lack the projections characteristic of the central pair apparatus (CPA). Mouse sperm doublets are not splayed, but they also have a pair of singlets extending beyond the axoneme **(Fig. 2e-f)**.

**Fig. 2.**
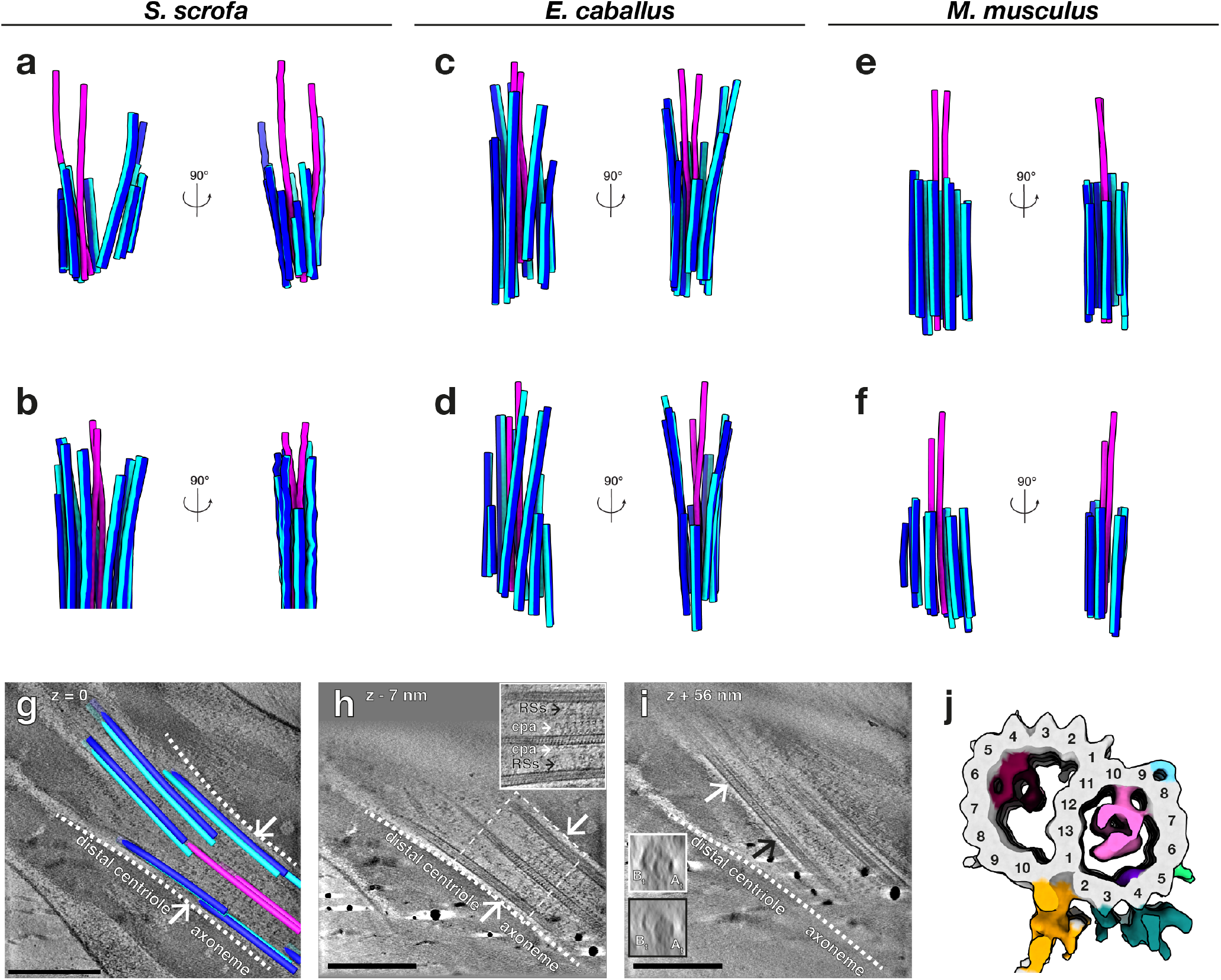
The distal centriole (DC) in mammalian sperm is composed of doublet microtubules arrayed asymmetrically around a pair of singlet microtubules. **(a-f)** Microtubules in the DC of pig **(a,b)**, horse **(c,d)**, and mouse **(e,f)** sperm traced from Volta phase plate cryo-tomograms of intact sperm. **(g-i)** Tomographic slices through cryo-FIB milled lamellae of the DC-to-axoneme transition in pig sperm show how the change in geometry **(g)** coincides with the appearance of axoneme accessory structures **(h)** and with density in the A-tubule **(i)**. **(j)***In situ* structure of the pig sperm DC with the tubulin backbone in grey and microtubule inner protein densities colored individually. **Labels:** RSs - radial spokes, cpa - central pair apparatus, A_t_ - A-tubule, Bt - B-tubule. **Scale bars:** 250 nm.

To more precisely define the DC-to-axoneme transition, we imaged cryo-FIB-milled sperm **(Fig. 2g-i)**. We directly observed this transition *in situ* in pig sperm, defined by the appearance of axonemal accessory proteins such as the radial spokes (RSs) and the projections of the CPA **(Fig. 2h)**. The onset of the axoneme coincides with a change in micro-tubule geometry **(Fig. 2g)**, suggesting that the splayed-open doublets are indeed characteristic of the DC. The transition zone also coincides with an increase in density in the A-tubule **(Fig. 2i)**, suggesting that binding of axonemal accessory structures is related to the regulated binding of A-tubule MIPs. We then determined the structure of DC doublets, revealing the presence of MIPs distinct from those in the PC **(Fig. 2j)**.

The flagellar waveform depends greatly on the properties of the base (Riedel-Kruse and Hilfinger, 2007), but there is very little information on how this region is organized in three dimensions in any cell type. In order to capture the full three-dimensional complexity of the flagellar base, we took advantage of enhanced contrast provided by the VPP, which allowed us to trace microtubules while retaining the context of the surrounding connecting piece **(Fig. 3a-c)**. Semi-automated neural-network-based segmentation (Chen et al., 2017) revealed that the connecting piece forms a large chamber enclosing the sperm centrioles. Although precise dimensions and shapes of the connecting piece differ across species **(Fig. 3d-f)**, its general architecture appears to be conserved across mammalian species.

**Fig. 3.**
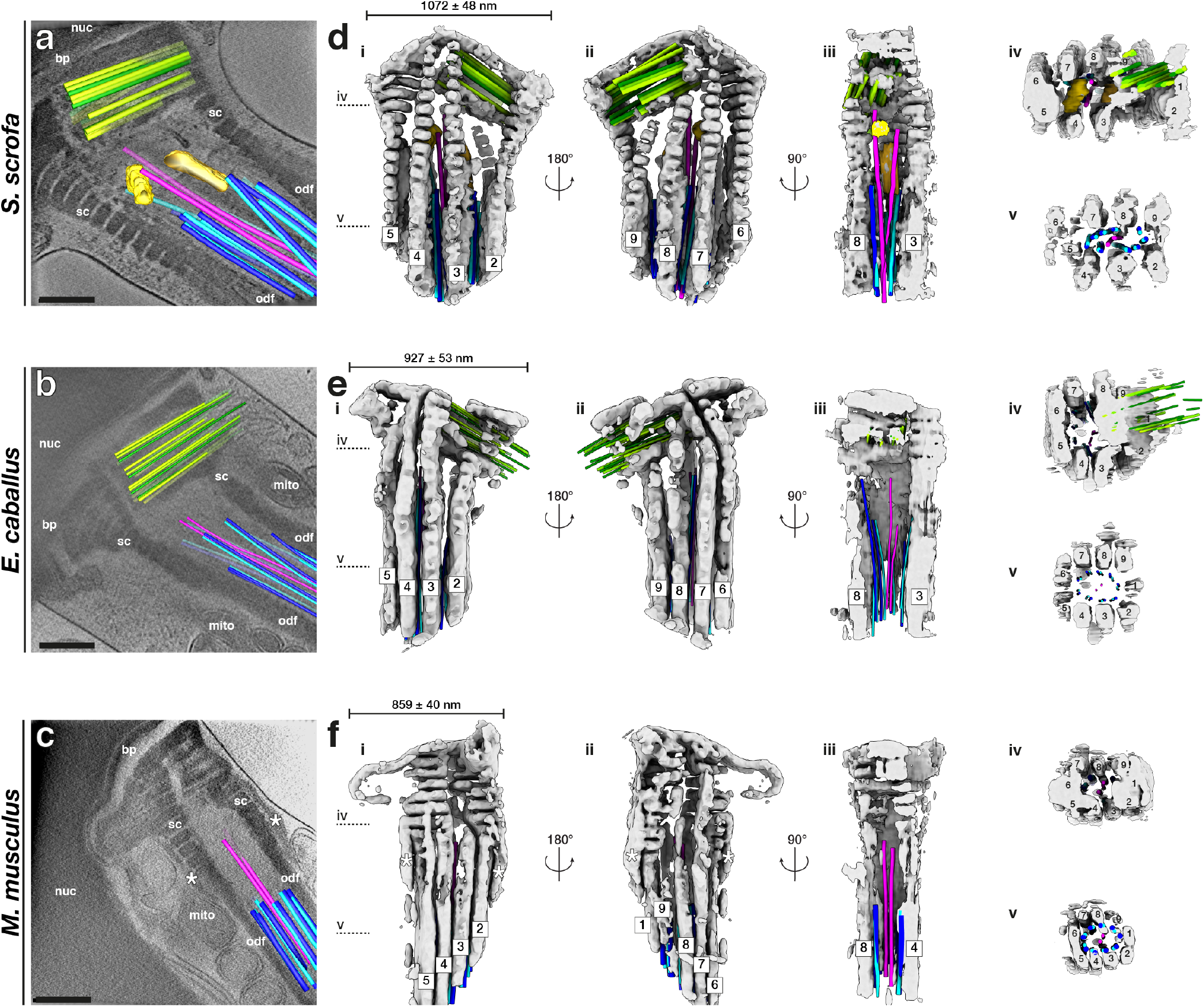
The connecting piece forms a large, asymmetric chamber around the sperm centrioles. **(a-c)** Slices through Volta phase plate cryo-tomograms of the neck region in intact pig **(a)**, horse **(b)**, and mouse **(c)** sperm. **(d-f)** Three-dimensional architecture of the flagellar base, with the connecting piece in grey, the proximal centriole in green, distal centriole doublets in blue and singlets in pink, and electron-dense bars in yellow. The connecting piece was segmented semi-automatically with a neural network, while microtubules were traced manually. **Labels:** nuc - nucleus, bp - baseplate, sc - striated columns, odf - outer dense fibers, mito - mitochondria. **Scale bars:** 250 nm.

The proximal region of the connecting piece consists of striated columns (SCs), called such because of their banded appearance. Following the numbering scheme laid out in (Ounjai et al., 2012), we found that the SCs follow a conserved pattern of grouping and splitting. The proximal connecting piece can be grossly divided into left and right regions. The right region forms the proximal centriolar vault where columns 8, 9, 1, 2, and 3 merge whereas the left region comprises columns 4, 5, 6, and 7 **(Fig. 3d-f, panels iv)**. The columns gradually separate, eventually splitting into nine separate columns that fuse distally with the ODFs **(Fig. 3d-f, panels v)**.

The connecting piece displays both marked left-right asymmetry and dorsoventral asymmetry in all three species. The PC is embedded within the proximal region of the connecting piece, and always on the same side. In pig sperm, one side of the proximal centriolar vault is formed by the Y-shaped SC 9, which also gives the entire connecting piece dorsoventral asymmetry **(Fig. 3d, panel ii)**. The material of the connecting piece extends through the interstices of the PC triplets **(Fig. 1a-c)** and is continuous with electron-dense material within the proximal lumen of the PC **(Fig. 1a’)**. Intriguingly, the dorsoventral asymmetry of the pig sperm PC is defined relative to the connecting piece, with the side of the shorter triplets always facing the Y-shaped segmented column 9 **(Fig. 3d)**.

The pig sperm connecting piece also has two electron-dense bars associated with the central singlets of the DC **(Fig. 3a,d, yellow and goldenrod)**, which resemble the bars observed in the bovine sperm connecting piece (Ounjai et al., 2012). These bars are conspicuously absent from horse and from mouse sperm. Instead, mouse sperm have two electron-dense structures flanking the SCs **(asterisks in Fig. 3c,f)**, an arrangement reminiscent of the distribution pattern of the centrosomal protein speriolin (Goto et al., 2010; Ito et al., 2019).

### The mammalian sperm axoneme anchors unique accessory structures and species-specific microtubule inner proteins

To gain insight into the molecular architecture of the axoneme, we determined *in situ* structures of the central pair apparatus **(Fig. 4)** and of the 96-nm axonemal doublet repeat **(Fig. 5)**. Our structures of the CPA are the first from any mammalian system, and our structures of the doublets are the first from any mammalian sperm, thus filling crucial gaps in the gallery of axoneme structures. The overall architecture of the mammalian CPA projection network is similar across the three species we examined **(Fig. S2)** and resembles that of the CPA from *Chlamydomonas* and from sea urchin sperm (Carbajal-González et al., 2013; Fu et al., 2019). Indeed, mutations in hydin, a component of the C2b projection, impair ciliary motility in both *Chlamydomonas* and mice (Lechtreck et al., 2008). However, the mammalian sperm CPA lacks the C1f projection found between the C1b and C1d projections in *Chlamydomonas* and sea urchin.

**Fig. 4.**
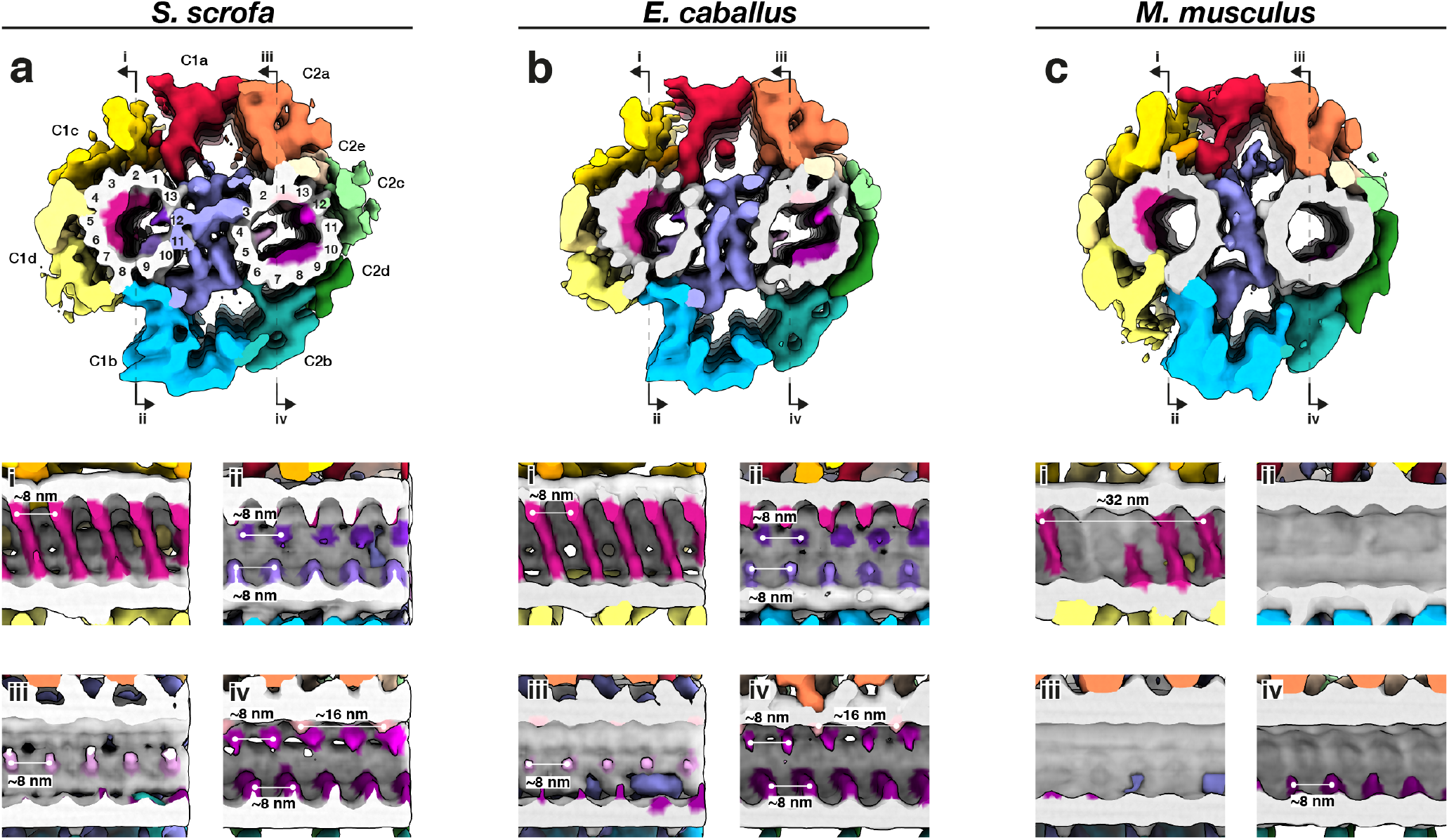
The mammalian sperm central pair apparatus (CPA) has a conserved projection network but species-specific micro-tubule inner proteins. **(a-c)** Whole-population *in situ* structures of the 32-nm CPA repeat from pig **(a)**, horse **(b)**, and mouse **(c)** sperm.

**Fig. 5.**
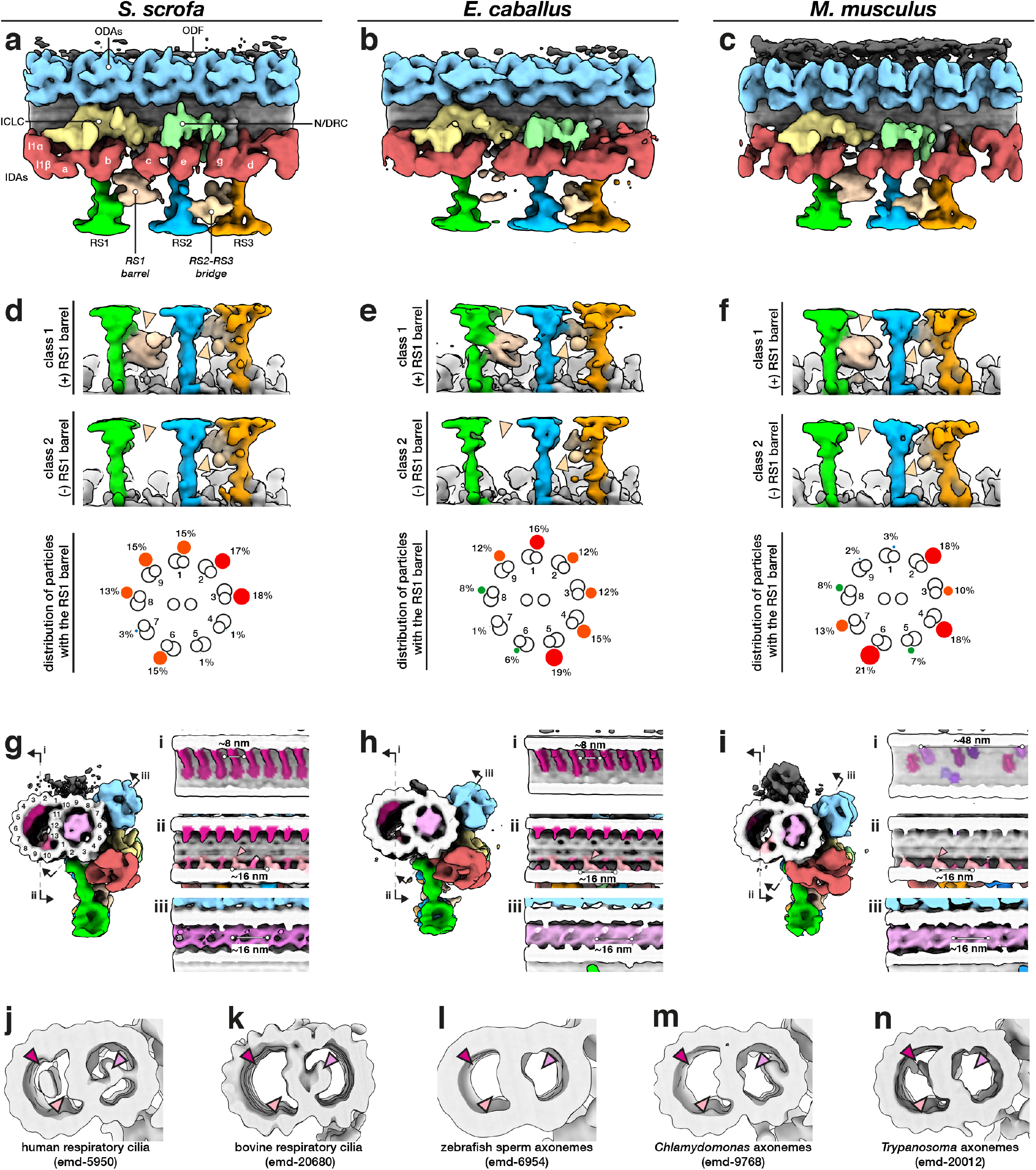
The mammalian sperm axoneme anchors unique accessory structures and species-specific microtubule inner proteins. **(a-c)** Whole-population *in situ* structures of the 96-nm repeat from pig **(a)**, horse **(b)**, and mouse **(c)** sperm. **(d-f)** Classification focused on the RS1 barrel revealed two distinct classes of particles, one with (top panels) and one without (middle panels) the structure. Particles with the RS1 barrel are distributed asymmetrically around the axoneme (bottom panels). **(g-i)** Microtubule inner proteins in axonemes from pig **(g)**, horse **(h)**, and mouse **(i)** sperm. **(j-n)** Microtubule inner proteins in axonemes from other cell types and organisms.

Mammalian CPAs have several MIPs **(Fig. 4)** that are absent from *Chlamydomonas* and from sea urchin sperm, which have only small MIPs or no MIPs respectively. Similar to the PC **(Fig. 1d)**, pig and horse CPAs have large helical MIPs in the C1 microtubule that bridge protofilaments 1-7 and repeat every 8 nm **(Fig. 4a-b, panels i)**. The mouse CPA has smaller MIPs in the same area, but these bridge fewer protofilaments and repeat with an overall periodicity of 32 nm **(Fig. 4c, panel i)**. Pig and horse also have smaller C1 MIPs on the side facing the bridge, with one bridging protofilaments 9 and 10 and the other jutting out from between protofilaments 12 and 13 **(Fig. 4a-b, panels ii)**. Both of these MIPs repeat every 8 nm and both are absent from the mouse **(Fig. 4c, panel ii)**. In the C2 microtubule, the pig and the horse have several other MIPs that are also absent from the mouse. These include a MIP that protrudes out from between protofilaments 4 and 5 and repeats every 8 nm **(Fig. 4a-b, panel iii)**, a MIP that binds between protofilaments 1 and 13 and repeats every 16 nm, and a MIP that extends from protofilament 12 and repeats every 8 nm **(Fig. 4a-b, panel iv)**. A fourth B-tubule MIP bridges protofilaments 7-9 in the pig and the horse, but this MIP is smaller and only bridges protofilaments 8 and 9 in the mouse **(Fig. 4a-c, panels iv)**.

Our *in situ* structures of the 96-nm axonemal doublet repeat from mammalian sperm revealed density for attachment to the outer dense fibers (ODFs) as well as novel structural features associated with the radial spokes (RSs) **(Fig. 5, Fig. S3)**. In particular, we observed a barrel-shaped structure associated with RS1 (the RS1 barrel) and an extensive bridge linking the stalks of RS2 and RS3 (the RS2-RS3 bridge). These structures are absent from all other axoneme structures reported so far **(Fig. S3a-g)**.

By focused classification, we resolved two distinct classes of particles, one with and the other without density for the RS1 barrel **(Fig. 5d-f)**. By separating out only particles with the barrel, classification also allowed us to improve the density for this structure. The RS1 barrel is ~18 nm long and ~16 nm wide and makes two major contacts with RS1, one at the base of the head and one at the middle of the stalk. Interestingly, the RS1 barrel is distributed asymmetrically around the axoneme. Although the RS1 barrel is not particularly enriched in any individual doublet position, it seems to occur less frequently in specific positions. In the pig, only 1% of the particles were found in each of doublets 4 and 5, while only 3% were in doublet 7. In the horse, only 1% were found in doublet 7. In the mouse, only 2% were found in doublet 9 and 3% in doublet 1.

Comparing mammalian sperm axonemes to those from other species reveals large variations in MIP densities **(Fig. 5g-n)**. Mammalian sperm have a large MIP that fills almost the entire lumen of the A-tubule (the A-MIP) **(Fig. 5g-i, bottom panels)**, which explains why the A-tubule appears dark in cross-section. The A-MIP makes extensive contacts with the A-tubule, including protofilaments A1-A3, A5-A6, and A8-A13, and has an overall periodicity of ~16 nm. Because the sperm A-MIP makes contacts with nearly all protofilaments of the A-tubule, it seems plausible that it would affect the mechanics of the doublet. The A-MIP also makes contacts with the same protofilament 9 to which the ODFs attach, which suggests that it may also functionalize the outer surface of the A-tubule for ODF docking. A-tubule MIPs are present in axonemes from human **(Fig. 5j)** (Lin et al., 2014a) and bovine **(Fig. 5k)** respiratory cilia (Greenan et al., 2020), but these are not as extensive as the A-MIP in mammalian sperm **(Fig. S3h)**. Zebrafish **(Fig. 5l)** (Yamaguchi et al., 2018) and sea urchin sperm (Lin et al., 2014b; Nicastro et al., 2011) do not have large MIPs in the A-tubule, nor do *Chlamydomonas* **(Fig. 5m)** (Nicastro et al., 2011; Owa et al., 2019) or *Trypanosoma* **(Fig. 5n)** (Imhof et al., 2019).

In all axoneme structures reported so far, B-tubules contain MIP3a and MIP3b, which bind to protofilaments B9 and B10 with staggered ~16-nm repeats **(Fig. 5g-i)**. However, in mammalian sperm, MIP3a has an additional density that links it to protofilament A13 **(Fig. 5g-i, panels ii, pink arrowheads; Fig. S3h)**. In pig and horse sperm, a helical MIP with an ~8-nm repeat bridges protofilaments B2-B7 **(Fig. 5g-i, panels i)**. Mouse sperm do not have this large MIP and instead have smaller MIPs that bind with an overall apparent periodicity of ~48-nm **(Fig. 5i, panel i)**. Large B-tubule MIPs have so far only been seen in human respiratory cilia **(Fig. 5j)** and in *Trypanosoma* (the ponticulus, **Fig. 5n**), but the morphometry of these MIPs differs from the helical MIPs in mammalian sperm.

A crucial comparison comes from the structure of axonemes from mouse respiratory cilia (Ueno et al., 2012). Mouse respiratory cilia lack the large A-MIP that is so prominent in mouse sperm, which points to cell-type specific differences in the MIP repertoire. Similarly, the RS1 barrel and the RS2-RS3 bridge **(Fig. 5)** are present in mammalian sperm, but not in human respiratory cilia or zebrafish sperm **(Fig. S3d,e)**. Because the radial spokes are key regulators of flagellar motility, as evidenced by the fact that radial spoke defects cause a number of ciliopathies (Lin et al., 2014a), the RS1 barrel and the RS2-RS3 bridge are likely to play a significant role in modulating sperm motility. As with many of the structures we report, defining why exactly they are needed in mammalian sperm would help us better understand the intricacies of flagellar organization.

### Outer dense fibers are directly coupled to axonemal doublets in the principal piece but not in the midpiece

To resolve how the ODFs associate with the microtubule doublets of the axoneme, we aligned and averaged particles from the principal piece, then classified them with a mask on the ODF-doublet attachment. Our structures reveal that, for doublets associated with ODFs, the ODFs are directly coupled to protofilament 9 of the A-tubule **(Fig. 6a-c)**. The ODF-doublet attachment consists of a pair of linkers spaced ~8 nm apart, with each pair spaced ~16 nm from the next, yielding an apparent overall periodicity of ~16 nm that is consistent across species. Direct ODF-microtubule coupling provides the elusive structural mechanism by which forces from axoneme bending can be transmitted through the ODFs to the base of the flagellum. As such, in the principal piece, the ODFs should be considered a part of the axoneme proper. This increases the effective diameter of the axoneme and also translates to an increase in bending moments (Lindemann, 1996; Lindemann and Lesich, 2016). However, the ODFs themselves are not of the same size, taper along the flagellum and terminate at different points, which leads to an anisotropic bending stiffness along the tail (Gadêlha and Gaffney, 2019; Lindemann, 1996).

**Fig. 6.**
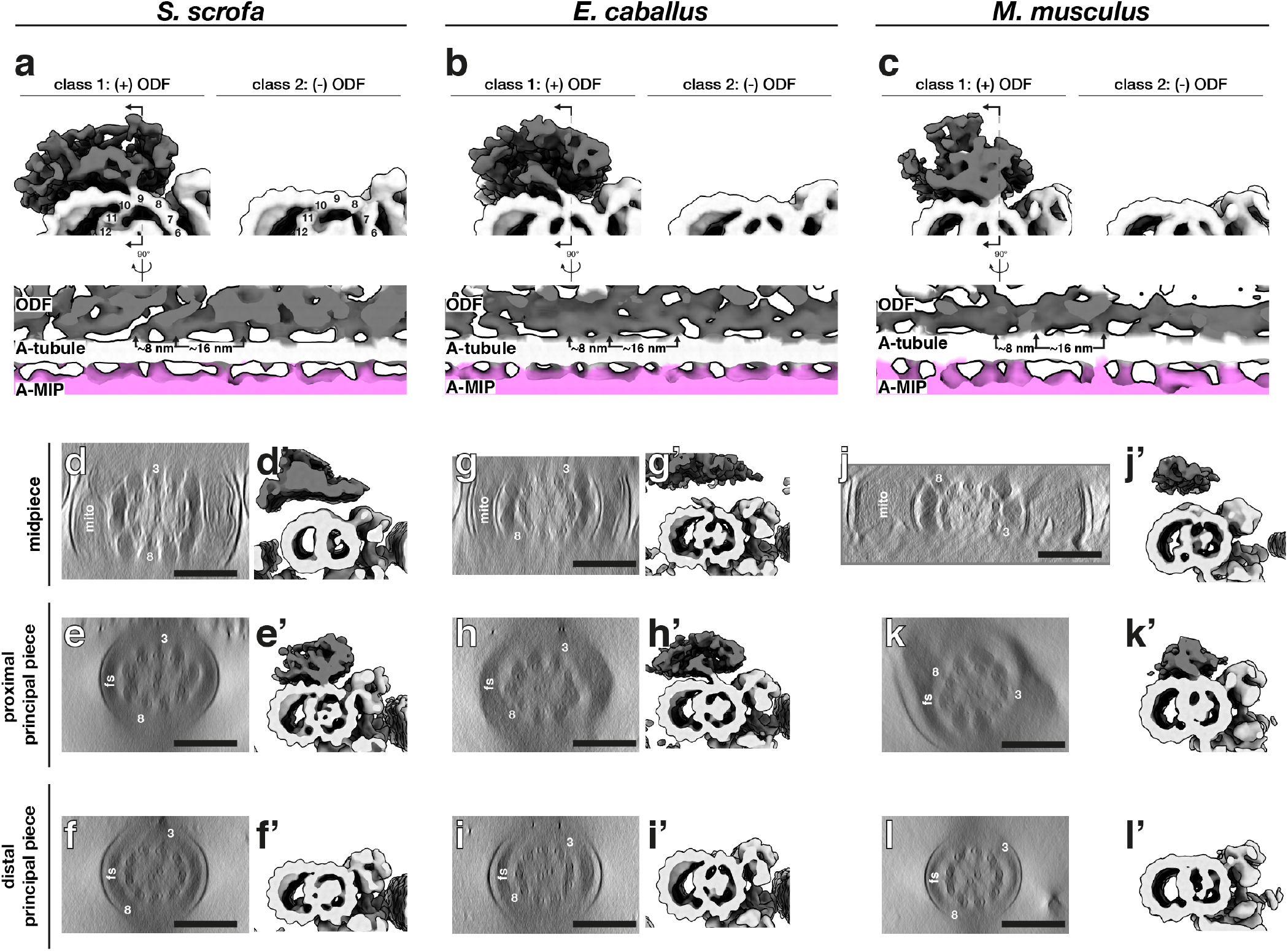
The attachment of outer dense fibers to axonemal doublets varies along the length of the sperm flagellum. **(a-c)** *In situ* structures of the 96-nm axonemal repeat from the principal piece of pig **(a)**, horse **(b)**, and mouse **(c)** sperm after classification focused on the ODF attachment. **(d-l)** Axoneme structures with particles from tomograms from **(d,g,j)** only the midpiece, **(e,h,k)** only the proximal principal piece, and **(f,i,l)** only the distal principal piece. **Labels:** ODF - outer dense fiber, mito - mitochondria, fs - fibrous sheath. **Scale bars:** 250 nm.

To determine how ODF-doublet association changes along the flagellum, we then separately averaged the 96-nm repeat from the midpiece, proximal principal piece, and distal principal piece **(Fig. 6d-l)**. Structures from the distal principal piece, after the ODFs had terminated, did not show density for the ODFs **(Fig. 6f’,i’,l’)**. In the proximal principal piece, the ODFs are directly attached to the axoneme via the A-tubule as described above **(Fig. 6e’,h’,k’)**. In the midpiece, the ODFs are at their largest but, surprisingly, are not directly connected to the microtubule doublets **(Fig. 6d’,g’,j’)**. This inhomogeneous pattern of association seems to be a general feature of mammalian sperm, as we observed it in all three species we examined. Our data thus reveal that the organization of accessory structures in mammalian sperm flagella is more complex than previously thought.

### Singlet microtubules in the endpiece are capped by a conserved plug but contain species-specific micro-tubule inner proteins

We then determined how the doublets of the axoneme transition into singlets of the endpiece **(Fig. 7a-c)**. We found that doublets can transition into singlets by two possible arrangements, either by a doublet splitting into two independent singlets **(Fig. 7d-f, left panels)** or by the B-tubule terminating abruptly **(Fig. 7d-f, right panels)**. Similar patterns have been observed in human sperm (Zabeo et al., 2019).

**Fig. 7.**
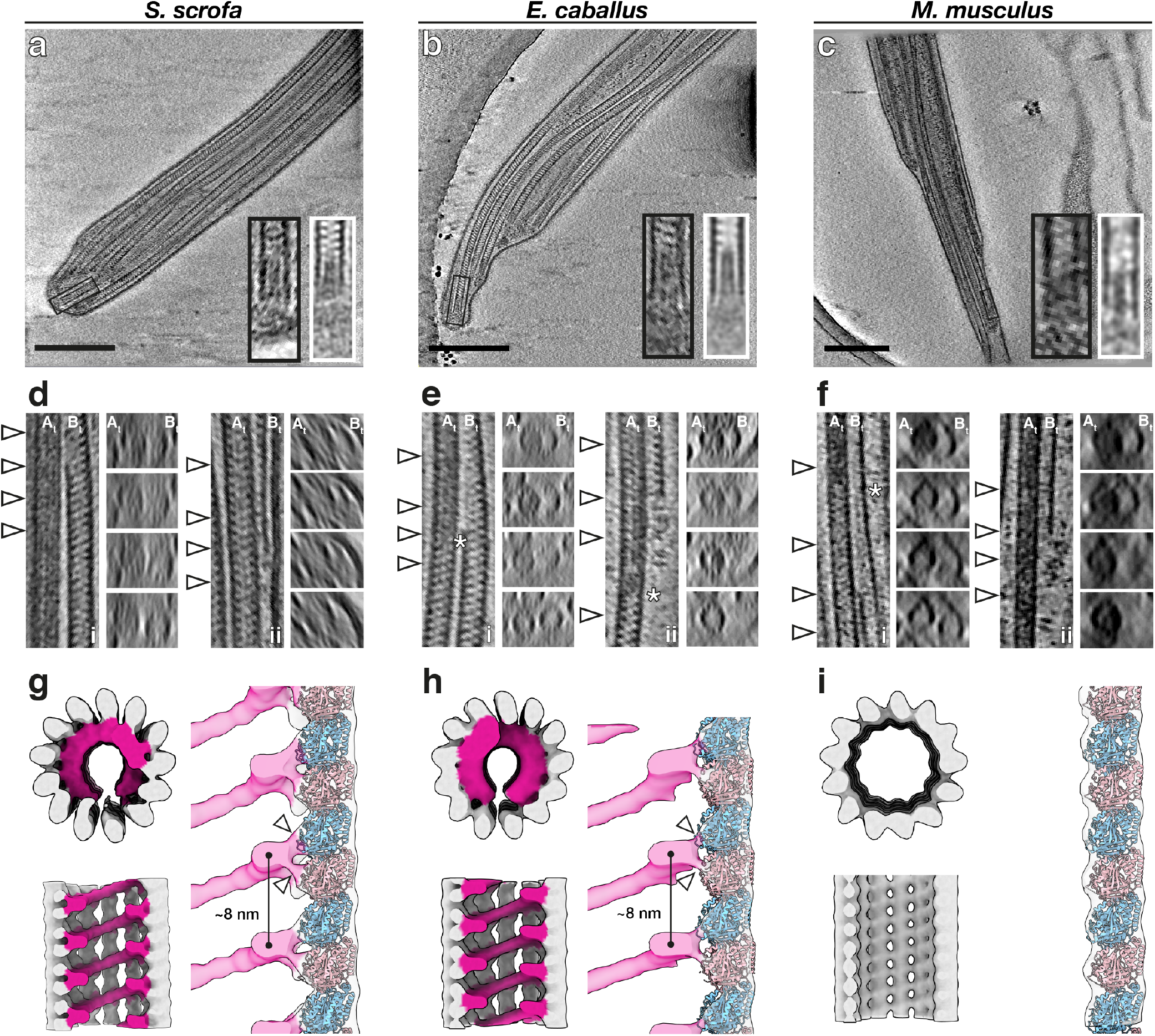
Singlet microtubules in the endpiece are capped by a conserved plug but contain species-specific microtubule inner proteins. **(a-c)** Slices through defocus cryo-tomograms of the endpiece in pig **(a)**, horse **(b)**, and mouse **(c)** sperm. **(d-f)** Representative examples of the mechanisms by which doublets can transition into singlets. Doublets can split into two complete singlets (panels i), or the B-tubule can terminate abruptly with the A-tubule extending as a singlet (panels ii). **(g-i)** *In situ* structures of singlet microtubules from pig **(g)**, horse **(h)**, and mouse **(i)** sperm endpieces. **Labels:** At - A-tubule, Bt - B-tubule. **Scale bars:** 250 nm.

We further observed that the doublet-to-singlet transition is also associated with loss of density in the A-tubule **(Fig. 7d-f, asterisks)**, although the precise location of this change relative to the splitting event varies. After the splitting event, 8-nm striations previously seen only in the B-tubule were visible in both singlets. The A-MIP thus seems to be a marker of the axoneme proper: the A-tubule lumen transitions from “empty” to “full” at the DC-to-axoneme transition **(Fig. 2i)**, whereas it goes from “full” to “empty” at the doublet-to-singlet transition.

We averaged endpiece singlets from the three species and found that they are consistently comprised of 13 protofilaments **(Fig. 7g-i)**. In the pig and the horse, singlets contain a helical MIP that follows the microtubule lattice, similar to the MIP previously described for human sperm endpieces (Zabeo et al., 2018). Our higher-resolution averages now reveal that this helical MIP makes independent contacts with both tubulin monomers **(Fig. 7g-h, arrowheads)**. This helical MIP is very similar to MIP7 we observed in the B-tubule of the PC **(Fig. 1d)**, to the helical MIP on the outer wall of the B-tubule of the axonemal doublets **(Fig. 5k-l)**, and to the helical MIPs in the CPA **(Fig. 4a-b)**. Intriguingly, this MIP is absent from endpiece singlets in the mouse **(Fig. 7i)**, where axonemal B-tubules and CPA microtubules also lack the large protofilament-bridging MIPs present in the pig and horse **(Fig. 4c, 4m)**.

We further observed that microtubule termini are capped in all three species **(Fig. 7a-c, black boxes)**. Averaging confirmed the presence of a plug extending 30 nm into the microtubule lumen **(Fig. 7a-c, white boxes)**. Normally, ciliary length control and turnover of axonemal components is mediated by the intraflagellar transport (IFT) system. However, IFT is absent in mature spermatozoa (San Agustin et al., 2015), which raises the question of how microtubule length and stability are maintained in these cells. These capping structures may stabilize free microtubule ends and prevent them from depolymerizing.

## Discussion

### Accessory structures are integral parts of mammalian sperm flagella and impose large mechanical asymmetries and anisotropies on the core axoneme

The presence of accessory structures in mammalian sperm flagella has long been recognized, but details of their three-dimensional organization and interactions with the core axoneme have remained elusive. Our comparative structural analysis revealed that the accessory elements impose striking multi-scale asymmetries and anisotropies on the sperm flagellum. Of particular relevance to wave generation, we found prominent asymmetry in the connecting piece at the base of the flagellum **(Fig. 3)**. This large-scale asymmetry could bias basal sliding to one side relative to the head, consequently polarizing inter-doublet sliding moments transmitted to the rest of the axoneme. Indeed, in order to swim forward, mouse sperm flagella must balance intrinsic asymmetry of the waveform with episodic switching of side-to-side asymmetric bends (Babcock et al., 2014). Asymmetric counterbends are also observed in rat sperm flagella, showing that shear developed in the distal flagellum is not identical when the flagellum is bent in the two opposing directions relative to the head (Lindemann et al., 2005). Polarized waveforms have also been reported for human and bovine sperm (Friedrich et al., 2010; Gadêlha et al., 2020; Saggiorato et al., 2017). Mathematical models incorporating basal sliding have confirmed the critical role of this phenomenon on waveform generation, but no modelling framework has attempted to study the effects of basal asymmetry that are now evident in our structures.

We also found that, in the principal piece, the ODFs are directly coupled to a defined protofilament on the axonemal doublets **(Fig. 5, 6)**. Such linkages provide a structural mechanism by which forces from axoneme bending can be transmitted through the ODFs to the base of the flagellum. Because the ODFs are anchored through the connecting piece at the base, these linkages also provide a mechanism by which changes in the base can be transmitted to the axoneme further down the tail. The ODFs also impose asymmetry and anisotropy on the flagellum; they gradually taper along the flagellum, but each ODF does so at a different rate, in contrast to the assumptions made in mathematical models incorporating the ODFs (Gadêlha and Gaffney, 2019; Riedel-Kruse and Hilfinger, 2007).

Beyond the gradual tapering and staggered termination of the ODFs, we show that a further source of proximal-distal asymmetry is the association of the ODFs with the axonemal doublets themselves **(Fig. 6)**. Intriguingly, this arrangement has actually been suggested previously based on measurements of ODF-doublet spacing in thin-section TEM of bull sperm (Lesich et al., 2014), and we now show that it holds true at the molecular level in three other mammalian species. This configuration would allow the ODFs to slide relative to the axoneme while being restrained by the mitochondrial sheath, lending flexibility to the midpiece. This is proposed to support formation of the extreme bends in the midpiece seen during hyperactivation (Lindemann and Lesich, 2016). Midpiece flexibility is crucial for sperm motility and hyperactivation, and mice lacking the catalytic subunit of a sperm-specific calcineurin isoform are infertile because they have more rigid midpieces (Miyata et al., 2015). Midpiece stiffness decreases as sperm transit through the epididymis (Jeulin et al., 1996; Miyata et al., 2015), which suggests that ODFs start out directly linked to the axoneme along their entire lengths and later locally detach in the midpiece.

Our comparative approach revealed that aforementioned details of how accessory structures relate to the microtubule core are conserved across species, suggesting that this architecture is fundamental for mammalian sperm motility. However, we also observe substantial inter-species variation, such as in the shape of the connecting piece **(Fig. 3)** and in the dimensions of ODFs **(Fig. 6)**. Comparative motility studies suggest that sperm flagella with larger ODFs form arcs with larger radii of curvature (Phillips, 1972). In contrast, how differences in the flagellar base and in head shape relate to differences in motility remains largely unexplored. We hope our structures will motivate and inform further theoretical and empirical work on the role of the base in shaping the flagellar beat.

### Mammalian sperm are characterized by large, helical, protofilament-bridging microtubule inner proteins that may affect microtubule bending stiffness

Our structures show that ciliary MIPs are highly diverse, both within a single cilium and across cell types **(Figs. 1, 2, 4, 5, 7)** By comparing several ciliary assemblies, we show that there are fundamental differences in the MIP repertoire across cell types and lineages. Most notably, large helical protofilament-bridging MIPs are present in essentially all microtubule-based assemblies throughout the sperm flagellum in pig and in horse, although they are reduced in corresponding structures in mouse sperm **(Figs. 4–5)**. These MIPs seem to be characteristic of mammalian sperm flagella as they are absent from axonemes of zebrafish sperm, sea urchin sperm, and mammalian respiratory cilia **(Fig. S3)**. Similar MIPs are present in human sperm endpiece singlets and in bovine respiratory epithelia, although in the latter they are restricted to the transition zone (Greenan et al., 2020). It is plausible that the helical MIPs are formed by variations of the same core protein complex, but direct confirmation awaits higher-resolution structures, genetic perturbation experiments, and direct labelling. If this were the case, however, this complex would have to adapt to the subtle differences in curvature between the walls of the centriolar/axonemal B-tubules and the 13-protofilament singlets.

The intimate and extensive contacts that the helical MIPs make with the tubulin lattice suggest that such a MIP would affect the mechanical properties of the microtubule itself. Microtubules are characterized by a paradoxical length-dependent bending stiffness attributed to the low shear modulus between adjacent protofilaments (Kurachi et al., 1995; Pampaloni et al., 2006; Taute et al., 2008). The protofilament-bridging MIPs observed here may decrease inter-protofilament shearing, increasing the shear modulus and the resulting bending stiffness of sperm microtubules. This adaptation might be necessary to withstand the large forces involved in moving the long flagellum of mammalian sperm, potentially reducing the length-dependency of bending stiffness in MIP-reinforced microtubules. These MIPs may also act as structural braces that suppress internal buckling within the axoneme under large loading in high viscosity fluids.

## Concluding Remarks

This study exemplifies the need for comparative studies of cilia and flagella, both from different species and from different cell types of the same species. Our study motivates future efforts to define how species-specific features of flagellar architecture affect the hydrodynamics of sperm motility. Such endeavours will likely involve the synthesis of structural cell biology, motility analysis, and mathematical modelling. The structures we present here provide crucial resources for understanding how the ancient and conserved ciliary microtubule core is ornamented to support motility through diverse fluid environments.

## Acknowledgements

The authors thank Dr. M Vanevic for enabling this project with superb computational support; Dr. D Vasishtan for providing scripts that greatly facilitated subtomogram averaging; Ingr. CTWM Schneijdenberg and JD Meeldijk for management and maintenance of the Utrecht University EM Square facility; Stal Schep (Tull en het Waal, The Netherlands) for providing horse semen; MW Haaker and M Houweling for providing mouse reproductive tracts; Prof. F Förster and Prof. A Akhmanova for critical reading of the manuscript; and Prof. EY Jones for insightful discussions. The authors also thank the Henriques Lab for the publicly-available 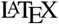 template. This work benefitted from access to the Netherlands Center for Electron Nanoscopy (NeCEN) with support from operators Dr. RS Dillard and Dr. CA Diebolder and IT support from B Alewijnse. This work was funded by NWO Start-Up Grant 740.018.007 to TZ, and MRL is supported by a Clarendon Fund-Nuffield Department of Medicine Prize Studentship.

## Author Contributions

PM, MZ, HH, EGB, and BMG provided sperm samples. MRL, MCR, and RTR prepared samples for cryo-EM. MRL performed cryo-FIB milling. MRL, MCR, RTR, SCH, and TZ collected cryo-ET data. MRL and MCR processed data. MRL, MCR, and TZ analyzed data. MRL, HB, and TZ wrote the manuscript, and all authors contributed to revisions.

## Materials and Methods

### Sperm collection and preparation

Pig sperm samples were purchased from an artificial insemination company (Varkens KI Nederland), stored at 18°C, and prepared for imaging within 1 day of delivery. Sperm were layered onto a discontinuous gradient consisting of 4 mL of 35% Percoll® (GE Healthcare) underlaid with 2 mL of 70% Percoll®, both in HEPES-buffered saline (HBS: 20 mM HEPES, 137 mM NaCl, 10 mM glucose, 2.5 mM KCl, 0.1% kanamycin, pH 7.6) and centrifuged at 750*g* for 15 min at RT (Harrison et al., 1993). Pelleted cells were washed once in phosphate-buffered saline (PBS: 137 mM NaCl, 3 mM KCl, 8 mM Na_2_HPO_4_, 1.5 mM KH_2_PO_4_, pH 7.4), resuspended in PBS and counted with a hemocytometer.

Horse semen was collected from mature Warmblood stallions using a Hanover artificial vagina in the presence of a teaser mare. After collection, semen was filtered through gauze to remove gel fraction and large debris, then transported to the laboratory at 37°C and kept at room temperature until further processing. Semen was diluted in INRA96 (IMV Technologies) to obtain a sperm concentration of 30 x 106 cells/mL. After this, sperm were centrifuged through a discontinuous Percoll gradient as described above for pig sperm for 10 min at 300*g* followed by 10 min at 750*g* (Harrison et al., 1993). The remaining pellet was resuspended in 1 mL of PBS and centrifuged again for 5 min at 750*g*.

Mouse sperm were collected from the cauda epididymis of adult male C75BL/6 mice as described in (Hutcheon et al., 2017). Briefly, male mice were culled as described in (Mederacke et al., 2015) and the cauda epididymides were dissected with the vas deferens attached and placed in a 500 μL droplet of modified Biggers, Whitten, and Whittingham media (BWW: 20 mM HEPES, 91.5 mM NaCl, 4.6 mM KCl, 1.7 mM D-glucose, 0.27 mM sodium pyruvate, 44 mM sodium lactate, 5 U/mL penicillin, and 5 μg/mL streptomycin, adjusted to pH 7.4 and an osmolarity of 300 mOsm/kg). To retrieve the mature cauda spermatozoa from the epididymides, forceps were used to first gently push the stored sperm from the vas deferens, after which two incisions were made with a razor blade in the cauda. Spermatozoa were allowed to swim out of the cauda into the BWW over a period of 15 min at 37°C, after which the tissue was removed and sperm were loaded onto a 27% Percoll density gradient and washed by centrifugation at 400*g* for 15 min. The pellet consisting of an enriched sperm population was resuspended in BWW and again centrifuged at 400*g* for 2 min to remove excess Percoll.

### Cryo-EM grid preparation

Typically, 3 μL of a suspension containing either 2-3 × 10^6^ cells/mL (for whole cell tomography) or 20-30 × 10^6^ cells/mL (for cryo-FIB milling) was pipetted onto a glow-discharged Quantifoil R 2/1 200-mesh holey carbon grid. One μL of a suspension of BSA-conjugated gold beads (Aurion) was added, and the grids then blotted manually from the back (opposite the side of cell deposition) for ~3 s (for whole cell tomography) or for ~5-6 s (for cryo-FIB milling) using a manual plunge-freezer (MPI Martinsreid). Grids were immediately plunged into a liquid ethane-propane mix (37% ethane) (Tivol et al., 2008) cooled to liquid nitrogen temperature. Grids were stored under liquid nitrogen until imaging.

### Cryo-focused ion beam milling

Grids were mounted into modified Autogrids (ThermoFisher) for mechanical support. Clipped grids were loaded into an Aquilos (ThermoFisher) dual-beam cryo-focused ion beam/scanning electron microscope (cryo-FIB/SEM). All SEM imaging was performed at 2 kV and 13 pA, whereas FIB imaging for targeting was performed at 30 kV and 10 pA. Milling was typically performed with a stage tilt of 18°, so lamellae were inclined 11° relative to the grid. Each lamella was milled in four stages: an initial rough mill at 1 nA beam current, an intermediate mill at 300 pA, a fine mill at 100 pA, and a polishing step at 30 pA. Lamellae were milled with the wedge pre-milling technique described in (Schaffer et al., 2017) and with expansion segments as described in (Wolff et al., 2019).

### Tilt series acquisition

Tilt series were acquired on either a Talos Arctica (ThermoFisher) operating at 200 kV or a Titan Krios (ThermoFisher) operating at 300 kV, both equipped with a post-column energy filter (Gatan) in zero-loss imaging mode with a 20-eV energy-selecting slit. All images were recorded on a K2 Summit direct electron detector (Gatan) in either counting or super-resolution mode with dose-fractionation. Tilt series were collected using SerialEM (Mastronarde, 2005) at a target defocus of between −4 and −6 μm (conventional defocus-contrast) or between −0.5 and −1.5 μm (for tilt series acquired with the Volta phase plate). Tilt series were typically recorded using either strict or grouped dose-symmetric schemes, either spanning ± 56° in 2° increments or ± 54° in 3° increments, with total dose limited to ~100 *e*^-^/Å^2^.

### Tomogram reconstruction

Frames were aligned either post-acquisition using Motioncor2 1.2.1 (Zheng et al., 2017) or on-the-fly using Warp (Tegunov and Cramer, 2019). Frames were usually collected in counting mode, but when appropriate super-resolution frames were binned 2X during motion correction. Tomograms were reconstructed in IMOD (Kremer et al., 1996) using weighted back-projection, with a SIRT-like filter (Zeng, 2012) applied for visualization and segmentation. Defocus-contrast tomograms were CTF-corrected in IMOD using *ctfphaseflip* while VPP tomograms were left uncorrected.

### Tomogram segmentation

Segmentation was generally performed semi-automatically using the neural network-based workflow implemented in the TomoSeg package in EMAN 2.21 (Chen et al., 2017). Microtubules, however, were traced manually in IMOD. Segmentation was then manually refined in Chimera 1.12 (Pettersen et al., 2004) or in ChimeraX (Goddard et al., 2018). Visualization was performed in ChimeraX.

### Subtomogram averaging

Subtomogram averaging with missing wedge compensation was performed using PEET 1.13.0 (Heumann et al., 2011; Nicastro et al., 2006). Resolution was estimated using the Fourier shell correlation (FSC) at a cut-off of 0.5 (Nicastro et al., 2006). Alignments were generally performed first on binned data, after which aligned positions and orientations were transferred to less-binned data using scripts generously provided by Dr. Daven Vasishtan. After alignment, classification was performed in order to assess heterogeneity and to identify cases of misalignment. Missing-wedge corrected classification was performed by first running a principal components analysis using the *pca* function, followed by k-means clustering using the *clusterPca* function (Heumann et al., 2011). The resulting class averages were manually inspected, and similar classes were combined. Specific averaging strategies are described below. Details of particle numbers and resolution estimates for each average are reported in **Table S1**.

### Proximal centriole triplets, axonemal doublets, central pair apparatus

Microtubule-based structures were manually traced in IMOD, and model points were added every 8 nm for PC triplets (Greenan et al., 2018), 32 nm for the CPA (Carbajal-González et al., 2013), or 96 nm for axonemal doublets (Nicastro et al., 2006) using *addModPts*. Subtomograms of approximately 70 nm x 42 nm x 70 nm (for centriole triplets), 100 nm x 100 nm x 100 nm (for the CPA), and 100 nm x 100 x 100 nm (for doublets) were computationally aligned and averaged.

For averaging triplets and doublets, particles with similar orientations (e.g; positions 9, 1, and 2) were first averaged. Sub-averages were then manually rotated along the y-axis using *modifyMotiveList* to align them with a common reference, followed by an alignment with a restricted search around the y-axis. If necessary, sub-averages were flipped to the right orientation using *modifyMotiveList* before a final restricted alignment in order to generate grand averages.

### Endpiece singlets

Endpiece singlets were manually traced in IMOD, and model points were added every 8 nm using *addModPts*. Subtomograms of approximately 35 nm x 35 nm x 35 nm were computationally aligned and averaged. An initial alignment was performed to align protofilaments, after which a mask was used to focus alignment on the helical MIP. The mask was then expanded to include the microtubule and, after classification, a final restricted alignment was performed.

---

**Fig. S1.**
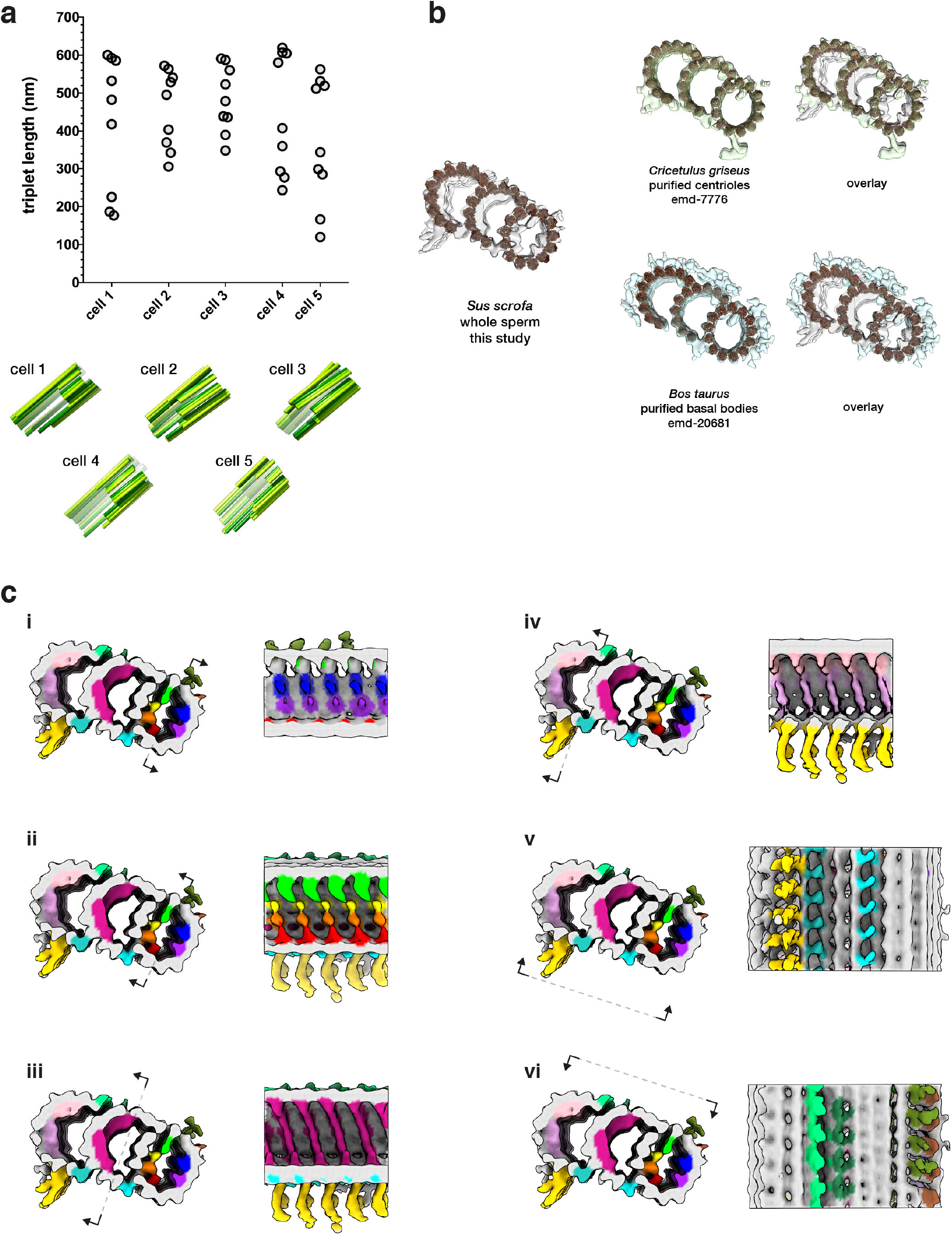
Structural features of the pig sperm proximal centriole (PC). **(a)** PC triplets have unequal lengths (top panel) and shorter triplets are grouped on one side, giving the PC dorsoventral asymmetry (bottom panel). **(b)** Many of the microtubule inner proteins (MIPs) in the pig sperm PC are not found in other mammalian centriole structures. **(c)** Details of the MIP densities in the pig sperm PC.

**Fig. S2.**
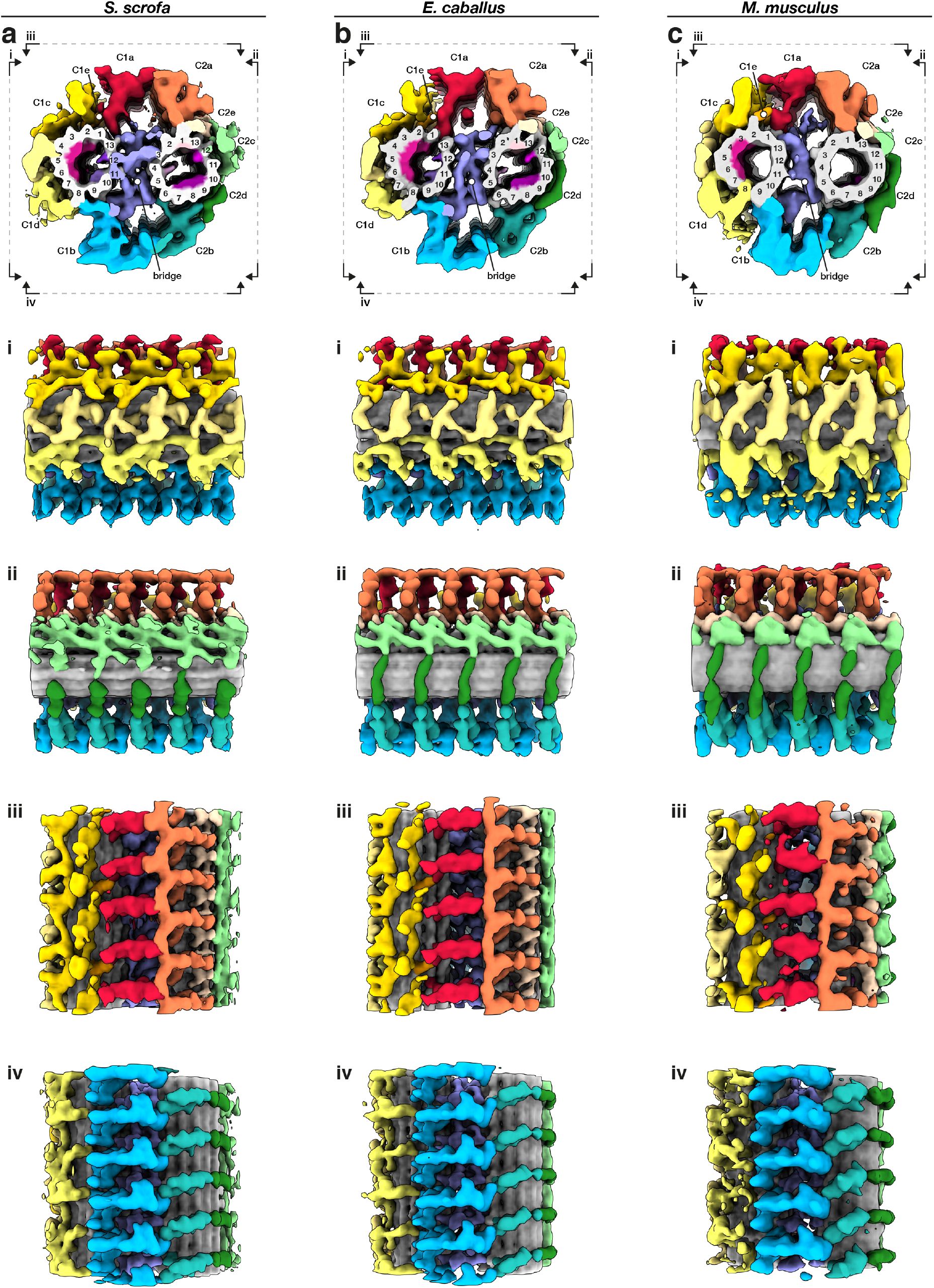
Structural details of the central pair apparatus projection network in mammalian sperm. **(a-c)** Whole-population *in situ* structures of the 32-nm CPA repeat from pig **(a)**, horse **(b)**, and mouse **(c)** sperm. Individual rotated views show details of C1 projections (i), C2 projections (ii), the interface between C1a and C2a projections (iii), and the interface between C1b and C2b projections (iv).

**Fig. S3.**
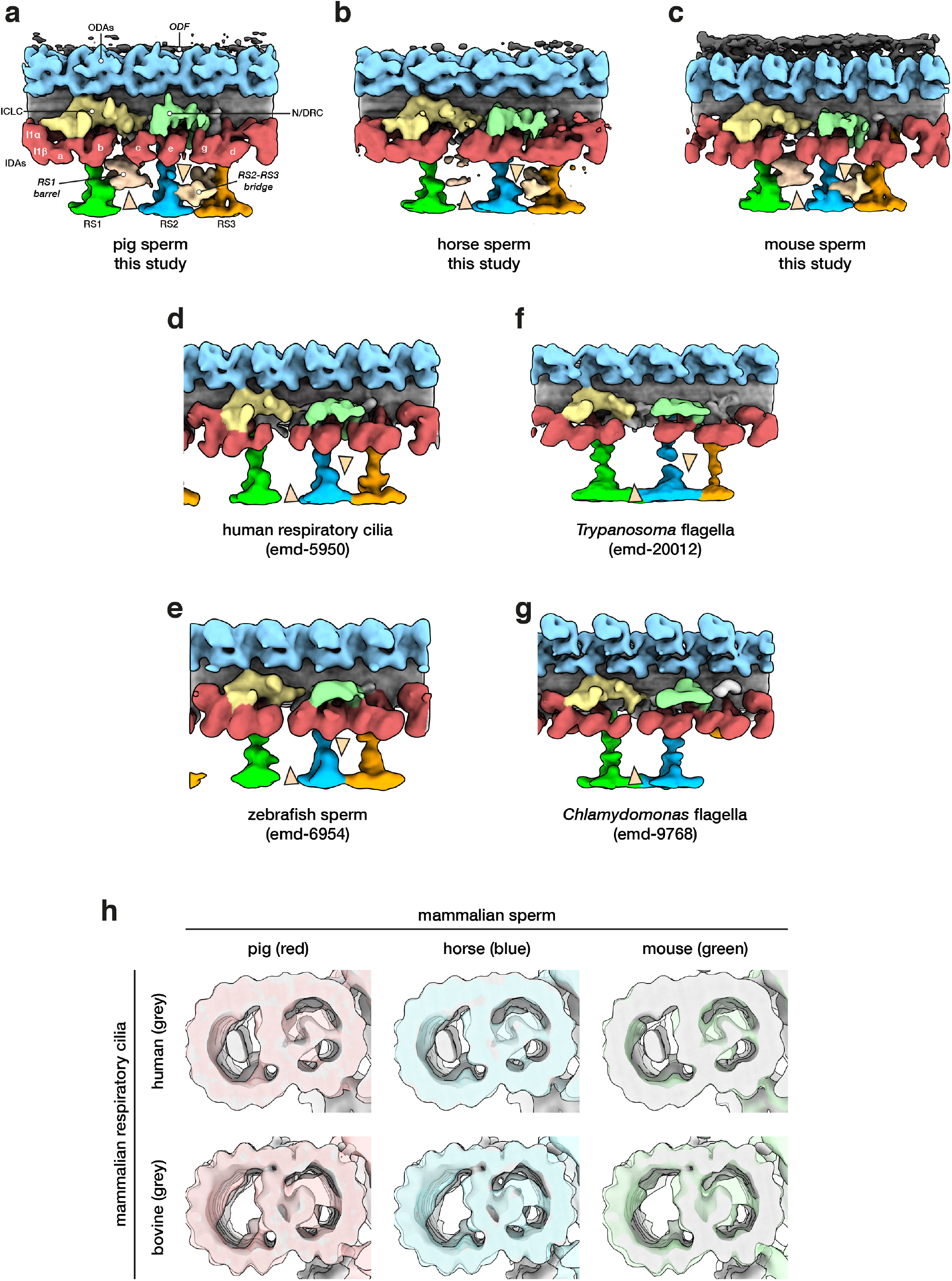
Comparing the mammalian sperm axoneme with structures from other cell types and species. **(a-g)** Structures of the 96-nm axonemal repeat from various species in cell types. Arrowheads mark the novel RS1 barrel and RS2-RS3 bridge so far seen only in mammalian sperm. **(h)** A-tubule MIPs are present in mammalian respiratory cilia, but the A-MIP in mammalian sperm is larger and makes more extensive connections with the A-tubule itself.

**Table S1.**
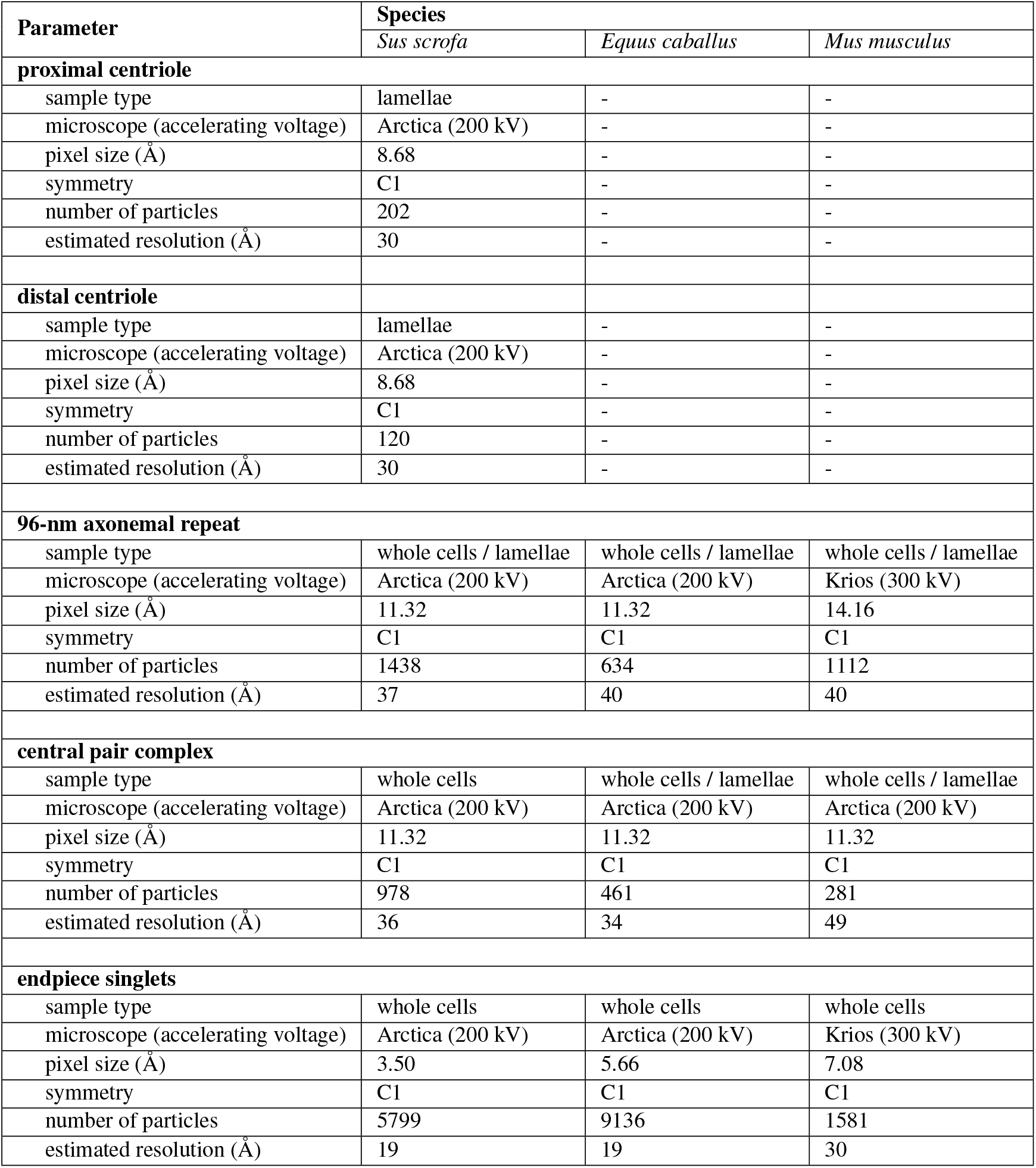
Summary of image acquisition parameters and processing metrics for subtomogram averaging.

| Parameter | Species |  |  |
| --- | --- | --- | --- |
|  | *Sus scrofa* | *Equus caballus* | *Mus musculus* |
| **proximal centriole** |  |  |  |
| sample type | lamellae | - | - |
| microscope (accelerating voltage) | Arctica (200 kV) | - | - |
| pixel size (Å) | 8.68 | - | - |
| symmetry | C1 | - | - |
| number of particles | 202 | - | - |
| estimated resolution (Å) | 30 | - | - |
| **distal centriole** |  |  |  |
| sample type | lamellae | - | - |
| microscope (accelerating voltage) | Arctica (200 kV) | - | - |
| pixel size (Å) | 8.68 | - | - |
| symmetry | C1 | - | - |
| number of particles | 120 | - | - |
| estimated resolution (Å) | 30 | - | - |
| **96-nm axonemal repeat** |  |  |  |
| sample type | whole cells / lamellae | whole cells / lamellae | whole cells / lamellae |
| microscope (accelerating voltage) | Arctica (200 kV) | Arctica (200 kV) | Krios (300 kV) |
| pixel size (Å) | 11.32 | 11.32 | 14.16 |
| symmetry | C1 | C1 | C1 |
| number of particles | 1438 | 634 | 1112 |
| estimated resolution (Å) | 37 | 40 | 40 |
| **central pair complex** |  |  |  |
| sample type | whole cells | whole cells / lamellae | whole cells / lamellae |
| microscope (accelerating voltage) | Arctica (200 kV) | Arctica (200 kV) | Arctica (200 kV) |
| pixel size (Å) | 11.32 | 11.32 | 11.32 |
| symmetry | C1 | C1 | C1 |
| number of particles | 978 | 461 | 281 |
| estimated resolution (Å) | 36 | 34 | 49 |
| **endpiece singlets** |  |  |  |
| sample type | whole cells | whole cells | whole cells |
| microscope (accelerating voltage) | Arctica (200 kV) | Arctica (200 kV) | Krios (300 kV) |
| pixel size (Å) | 3.50 | 5.66 | 7.08 |
| symmetry | C1 | C1 | C1 |
| number of particles | 5799 | 9136 | 1581 |
| estimated resolution (Å) | 19 | 19 | 30 |

